# Reach and Grasp Altered in Pantomime String-Pulling: A Test of the Action/Perception Theory in a Bilateral Reaching Task

**DOI:** 10.1101/679811

**Authors:** Surjeet Singh, Alexei Mandziak, Kalob Barr, Ashley A Blackwell, Majid H Mohajerani, Douglas G Wallace, Ian Q Whishaw

**Affiliations:** Department of Neuroscience, Canadian Centre for Behavioural Neuroscience (CCBN), University of Lethbridge, Lethbridge, Alberta, Canada; Department of Psychology, Northern Illinois University, De Kalb, Illinois, 60115 USA

**Author notes:** **Correspondence to**: Surjeet Singh, Department of Neuroscience, Canadian Centre for Behavioural Neuroscience (CCBN), University of Lethbridge, 4401 University Drive, Lethbridge, Alberta CA, T1K 3M4.

## Abstract

The action/perception theory of cortical organization is supported by the finding that pantomime hand movements of reaching and grasping are different from real movements. Frame-by-frame video analysis and MATLAB^®^ based tracking examined real/pantomime differences in a bilaterally movement, string-pulling, pulling down a rope with hand-over-hand movements. Sensory control of string-pulling varied from visually-direct when cued, visually-indirect when non cued and somatosensory controlled in the absence of vision. Cued grasping points were visual tracked and the pupils dilated in anticipation of the grasp, but when noncued, visual tracking and pupil responses were absent. In real string-pulling, grasping and releasing the string featured an arpeggio movement in which the fingers close and open in the sequence 5 through 1 (pinki first, thumb last); in pantomime, finger order was reversed, 1 through 5. In real string-pulling, the hand is fully opened and closed to grasp and release; in pantomime, hand opening was attenuated and featured a gradual opening centered on the grasp. The temporal structure of arm movements in real string-pulling featured up-arm movements that were faster than down-arm movement. In pantomime, up/down movements had similar speed. In real string-pulling, up/down arm movements were direct and symmetric; in pantomime, they were more circular and asymmetric. That pantomime string-pulling featured less motoric and temporal complexity than real string-pulling is discussed in relation to the action/perception theory and in relation to the idea that pantomimed string-pulling may feature the substitution of gestures for real movement.

**Significant Statement:** Most laboratory studies investigating hand movements made by humans feature single hand movements, the current study presents a novel string-pulling task to study bimanual coordination of left and right hands in real and pantomime conditions. The results show that pantomime string-pulling featured less motoric and temporal complexity than real string-pulling. These findings are relevant to the contemporary theory of action and perception that the dorsal stream (parietal cortex) is related to actions and the ventral stream (temporal cortex) is related to perception.

## Introduction

The investigation of hand movements is an area of central interest in the study of motor control, brain organization, robotics and evolution (Debaere et al., 2003; Karl et al., 2012; Karl and Whishaw, 2013; Kilteni and Ehrsson, 2017; Salvietti, 2018; Isa, 2019; Lemon, 2019; Wu et al., 2019). The action/perception theory suggests that there are two neural systems of movement control, an online action-system for real movements and an offline perceptual-system for pantomime movements, mediated by a dorsal stream parietal cortex to motor cortex pathway and by a ventral stream, temporal cortex to motor cortex pathway, respectively (Goodale et al., 1991; Goodale et al., 1994; Westwood et al., 2000; Milner et al., 2001; Fukui and Inui, 2013; Holmes et al., 2013; Kuntz and Whishaw, 2016). One focus of hand studies is on single hand movements, picking up an object, retrieving an item of food for eating, or pointing (Karl et al., 2012; Karl and Whishaw, 2013; Freud et al., 2018; Ingram et al., 2019; Urbán et al., 2019), another is on bilateral hand movements (Kelso et al., 1979; Franz et al., 1991; Donchin et al., 1998; Swinnen, 2002; De Jesus et al., 2018; Osumi et al., 2019; Shih et al., 2019), but real/pantomime differences have not been featured in bilateral hand movement studies. Because bilateral hand movements can be target directed and involve interlimb coordination, their complexity provides a dimension of movement that could provide insights into real/pantomime nervous system differences. The present study examines a bilateral reaching-to-grasp task, string-pulling.

String-pulling is a bilateral hand movement task requiring a string to be reeled to obtain a reward attached to its end. Jacobs and Osvath (Jacobs and Osvath, 2015) report that over 160 bee, bird, and mammalian species display versions of the behavior. The utility of the task is in assessing perceptual, learning and motor ability and cognitive functions, such as reasoning, insight learning, and social learning. One string pulling paradigm requires pulling a vertically oriented string with up/down, alternating hand movements (Blackwell et al., 2017; Blackwell et al., 2018a; Blackwell et al., 2018b). When a participant adopts an erect posture, facing a camera, the task provides an unobstructed view the torso, arm, hand and finger from which the coordination of the movements of both hands can be gauged. Humans engage in many string-pulling types of behavior (see also, (Taylor et al., 2012; Rat-Fischer et al., 2014)) and string-pulling tasks have been given to human infants and children (Richardson, 1932; Piaget, 1952; Redshaw, 1978; Brown, 1990; Silva et al., 2005; Silva et al., 2008; Albiach-Serrano et al., 2012), but there has been no description of adult human string pulling and no studies of pantomime string-pulling.

The first objective of the present study was to describe the arm, hand and finger movements and the sensory control of string pulling in human female and male participants. Variations of the task included conditions in which participants were instructed to use visual cues, freely pulled, or were blindfolded. The second objective of the study was to analyze the kinematics involved in the reach/grasp movements and the timing associated with the bilateral control of string-pulling. Participants wore color coded gloves on the left and right hand and their movements were tracked using MATLAB based software with a graphical user interface (Singh et al., 2016). The third objective of the study was to compare the movements of real string-pulling with pantomime string-pulling. The comparison of real and pantomime movements has been used to obtain insights in the cortical processes involved in online real actions vs perceptually based pretend actions. Male and female undergraduates were filmed from a frontal view and pulled a string (a soft rope), that was threaded through an overhead pulley, using hand-over-hand movements. Different groups of participants were tested in a number of conditions and after real string-pulling trials, the same participants were asked to pantomime the pulling movement that they had just made.

## Methods

### Participants

Participants were 68 undergraduate students aged 18 to 23 enrolled in a neuroscience class at the University of Lethbridge. Ion one experiment, 48 participants were divided into 4 groups of 12, with each group comprised of 6 females and 6 males. All participants were selected from a larger group of students on the basis of a short questionnaire that ascertained that they were right-handed for activities such as writing and throwing a ball. In a second experiment, 20 participants were divided into two groups of 10, and these participants performed string-pulling while wearing eye-tracking glasses. Before beginning the experiment, the participants read a short summary of the experiment, gave their written consent to participate, and gave consent to their images being used for experimental analysis and dissemination. The experiments were approved by the University of Lethbridge human subject’s ethics committee.

### String

Two 14 mm diameter strings (ropes) were used, each 1,341 cm long. One was a plain white string and the other had a 2 cm wide bands of black electrician’s tape wrapped around it to demark the string into 30 cm segments. The strings were suspended through a pully in the ceiling. A knot tied at the end of each string prevented the string from passing through the pulley. A smaller string tied to the knot allowed the main string to be pulled back to its starting position.

### Attire

Participants were asked to dress in a dark top and bottom. Before initiating string-pulling, the participants put on VWR Nitrile gloves (VRW International, LLC Ranor, PA). Participants put a blue glove on the right hand and a green glove on the left hand. Preliminary experiments in which participants wore gloves indicated that string pulling movements were similar in ungloved and gloved participants.

### Eye and Pupil Tracking and Visual Attention

#### Eye tracking and pupil measurement

To evaluate eye movements and pupil changes, participants pulled the real string and then pantomimed string-pulling while wearing eye-tracking glasses. Eye movements and the pupil were monitored using a Pupil-Lab eye tracker (Kassner et al., 2014). Pupil-Lab is a wearable mobile eye tracking headset with one scene camera and two infrared (IR) spectrum eye camera for dark pupil detection. The cameras connect to a laptop computer platform via high speed USB 2.0. The camera video streams were read using Pupil Capture software for real-time pupil detection and gaze mapping. Pupil captures a view of the gaze scene at 30 Hz and gaze focus and pupil diameter at 120 Hz. The World window of the software was used to display the video stream from the scene camera with gaze location superimposed on the scene as a small round dot. The gaze camera was calibrated using a 9-point calibration method and the accuracy of calibration was checked by having participants fixate their gaze on selected objects in the test room before and after string pulling. Frame-by-frame inspection of the World view video was used to determine whether participants were looking at the string when pulling and for determining what portion of the string was the focus of gaze. On line changes in pupil diameter were similarly monitored in relation to string-pulling hand movements.

#### Visual attention

In an experiment in which participants did not wear eye tracking glasses, the relationship between looking and reaching was gauged from their head orientation and grasp location in the y-plane. Orientation was assessed using a three-point scale: 1 = looking up, 0 = looking straight ahead, −1 = looking down) and the grasp location of the hand was also assessed on a three-point scale: 1 = reaching above the head, 0 = reaching to the level of the face, −1 = reaching below the level of the face. Correlations between head orientation and grasp location determined whether participants were looking toward the location on which their hand grasped the string. Head orientation and hand placement were rated for 4 up-down sequences by each hand from one real and one pantomime string pulling act to obtain a mean score.

### Release and Grasp Movements

A rating scale was used to quantify how the hands released and grasped the string. The order of finger extension when the hand was opening and finger flexion when the hand was closing was rated on a three-point scale. If the fingers extended or flexed in the order 5 through 1 (pinki first, thumb last) the movement was rated as “0”; if the fingers opened or closed concurrently, the movement was rated as “0.5”; if the fingers opened or closed in the order 5 through 1, the movement was rated as “1”. Five hand release and hand grasp movements were rated for each hand for each participant from one real and one pantomime string pulling act to obtain a mean score.

### Tracking Hand Movements

In computer vision, image segmentation is the process to cluster pixels into salient groups to identify regions of interest. Multiple methods exist to perform image segmentation e.g. thresholding methods Otsu’s (Otsu, 1979), color-based segmentation such as K-means clustering (Arthur and Vassilvitskii, 2007), transform methods such as watershed segmentation (Meyer, 1994) and texture filter methods (Gonzalez et al., 2003). Here participants wore green (left hand) and blue (right hand) nitrile gloves to provide high target contrast relative to the black background (body and wall sheet), thus, making image segmentation process amenable to a thresholding-based segmentation method. The Color Thresholder app in MATLAB was used to perform color-based segmentation of the left/right hand for each participant.

First, a random frame from the recorded behavior was opened in Color Thresholder app, which in turn displayed the image along with point clouds representing the same image in several popular color spaces: RGB, HSV, YCbCr, and L*a*b*. The color space that provided best color separation for left/right hands from the background was selected. The colorspace and range for each channel of the colorspace were set within the app and a thresholding function was autogenerated to recreate the segmentation in all frames of the video. Using a similar approach presented in (Singh et al., 2016), the centroid of hands was detected in each segmented frame and presented as the track plot once the video was processed.

To describe the movements of one hand with respect to another we extracted the instantaneous phase of the real-valued y-axis motion *y*(*t*) using an analytic signal representation as follows:

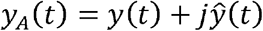

where, *y*_*A*_(*t*) is the analytic signal, 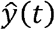 is the Hilbert transform of *y*(*t*). The analytic signal obtained is express it in exponential notation:

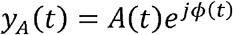

where, *A*(*t*) is the instantaneous amplitude and *ϕ*(*t*) is the instantaneous phase. *ϕ*(*t*) values gave us the timepoints when the participants were grasping or releasing the string.

The best fit to an ellipse for the given set of x-y coordinates of each hand’s trajectory (Bookstein, 1979; Gander et al., 1994) was used to describe the spatial occupancy patterns of left/right hand. A Least-Squares criterion for estimation was performed for the for the following conic representation:

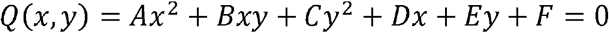

After finding the x-y coordinates of the center, length of long and short was used to find the area of the ellipse from which to extract frequency, amplitude, and the spatial occupancy patterns in real and pantomime tasks.

The MATLAB tracking procedure was used to assess one real and one string pulling act by each of the participants. In addition, a second string pulling act was manually digitized using PhysicsTracker (Open Source Physics (OSP) Java framework) to confirm the accuracy to the MATLAB procedure. Because the two methods gave the same statistical results only the MATLAB obtained results are reported here.

## Procedure

A participant was asked to stand in front of the string in front of a black sheet and at a fixed distance from the camera. Participants were asked to take the string in one hand to stabilize it. They were then instructed that on a “go” command they should pull the string with alternating left-and-right hand overhand pulls until the string was stopped by its knot. Participants were given no instructions with respect to which hand they should start with, but most participants used their right hand to stabilize the string. Each participant was given four trials:

1. *Practice*. The first trial was used as practice, and the participants pulled a real string to ensure that they understood the instructions.
2. *Real string-pulling*. Participants were given three trials in which they pulled the real string to its end.
3. *Pantomime*. The participants were given three trials in which they pantomimed string-pulling in the same way that they had pulled in the real string-pulling condition. For the pantomime trial, the string was moved to one side and the participant was instructed: “Now we would like you to pretend to pull the sting in the same way that you actually pulled it.” Each string-pulling trial generated about 4 to 7 pulling cycles by each hand.

### Experiment 1: condition comparisons

Twelve participants (6 female and 6 male) were assigned to each of the following groups:

1. *Control, unmarked string*. The control group pulled the plain string that had no markings.
2. *Control, marked string*. The control-marked group pulled a string that had black markings, but they were given no instructions with respect to the markings.
3. *Visual occluded*. The occluded group pulled the plain string that had no markings.
4. *Visual cue*. The visual cue group were asked to grasp the string at every second black cue on each pull with the left hand and on each pull with the right hand.

### Experiment 2: eye and pupil tracking

Participants who wore eye tracking glasses were assigned to each of the following groups:

1. *Control, marked string*. The control marked group pulled a string that had black markings, but they were given no instructions with respect to the markings. After performing two real string-pulling trials they performed two pantomime string-pulling trials.
2. *Visual cue*. The visual cue group were asked to grasp the string at every second black cue for each pull with the left hand and each pull with the right hand. After performing two real string-pulling trials they performed two pantomime string-pulling trials.

### Statistical analyses

The results were analyzed using repeated Analysis of Variance (ANOVA) for repeated measures and Bonferroni follow-up tests according to the IBM SPSS®. The independent variables were Groups (participant groups), Hands (left and right hands) Sex (male and female), and Conditions (real and pantomime string-pulling). Independent measures included Direction (up and down hand movements), Velocity (distance /sec) including Maximum and Minimum velocity of up and down movements, Frequency (number of pulls by a hand per/sec), Amplitude (distance/cm of up and down movements), Spatial Occupancy (the area enclosed by an up or down string pulling movement relative to a direct path), the Cartesian volume enclosed by a complete up/down hand excursion, and Asymmetry (difference in measure by one hand compared to the other). Pearson produce correlations were used to related gaze orientation to hand location. A value of p<0.05 was accepted as significant.

## Results

### Arm and hand movements

Figure 1 illustrates string-pulling movements in a representative participant, with a video of a real string-pull shown in Video 1 and pantomime sting-pulling shown in Video 2. String-pulling is a cyclic movement in which one forelimb alternates with the other to advance the string, with both forelimbs featuring characteristic upper arm, lower arm, and hand movements in each phase of the action. Table 1 summarizes the component acts of string-pulling. The movements are identifiably similar in both real and pantomime string-pulling and consist of the following acts:

1. *Release.* Release of the string by the pulling hand occurs by extension of the lower arm, upper arm, and the fingers to release the string.
2. *Lift.* The hand is lifted to the midpoint of the torso to initiate a new pulling movement by flexion of the lower arm at the elbow, with the hand carried to a horizontal position with the digits slightly flexed, and with the distal ends of the digits aligned to the body midline.
3. *Advance.* The hand is carried in an upward motion toward the string, first by extension of the upper arm at the shoulder followed by extension of the lower arm at the elbow, with a slight adduction of the hand so that palm of the hand is aligned with the body midpoint and with the string.
4. *Grasp.* To grasp the fingers more fully extend and open to purchase the string in an arpeggio movement in which the digits close in the sequence 5 through 1 for real string pulling or 1 through 5, and with contact with the string made the hand begins to descend.
5. *Pull.* For the pull, the upper arm flexes at the shoulder, the lower arm flexes at the elbow, and the fingers are fully closed on the string and the hand is dorsiflexed at the wrist, with the movements advancing the string toward the midpoint of the torso.

1. *Push.* For the push, the hand directs the string downward toward the hips by extension of the lower arm at the elbow and by pronation of the hand such that the digits are directed downward.

**Table 1:** Main Arm and Hand Movements of Real String Pulling

| Movement | Hand | Lower Arm | Upper Arm |
| --- | --- | --- | --- |
| Release extend | Fingers extend/open<br>Arpeggio 5-1 | Abduct , extend | Abduct |
| Lift extend | Fingers partially closed | Flex at elbow | May |
| Advance | Fingers extend | Extend at elbow | Extend |
| Grasp flex | Fingers flex/close Arpeggio<br>5-1 | Begins to close | Begins to |
| Pull | Fingers grasp | Flex at elbow | Flex |
| Push abduct | Fingers grasp | Extend at elbow | Flex |

**Figure 1.**
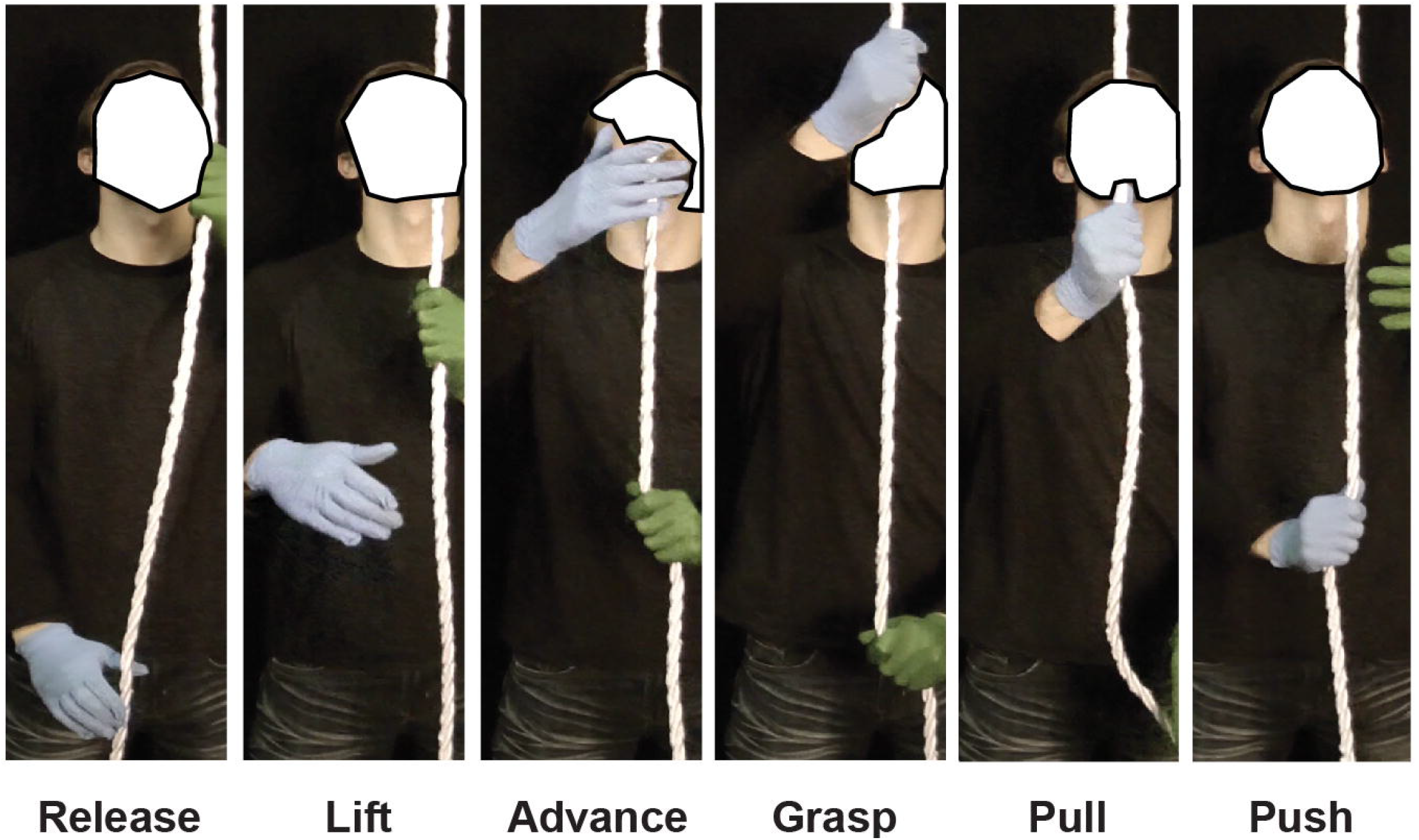
The five movements of string-pulling illustrated for the right hand of a participant making a real string-pulling movement. *Release*, fingers open and extend to release the string; *Lift*, the hand is raised to the midpoint of the torso largely by flexion of the lower arm; *Advance*, the hand is raise to grasp the string largely by extension of at the shoulder and elbow; *Grasp*, the fingers are closed and flexed to grasp the string; Pull, the string is lowered largely by flexion at the shoulder and the lower arm; *Push*, the continued movement of the string is produced largely be extension of the lower arm. Note: gaze is directed to the string at a point above the grasp point on the string.

### Eye and Pupil Tracking and Visual Attention

Figure 2 illustrates the main findings related to eye, pupil and visual attention for string-pulling.

**Figure 2.**
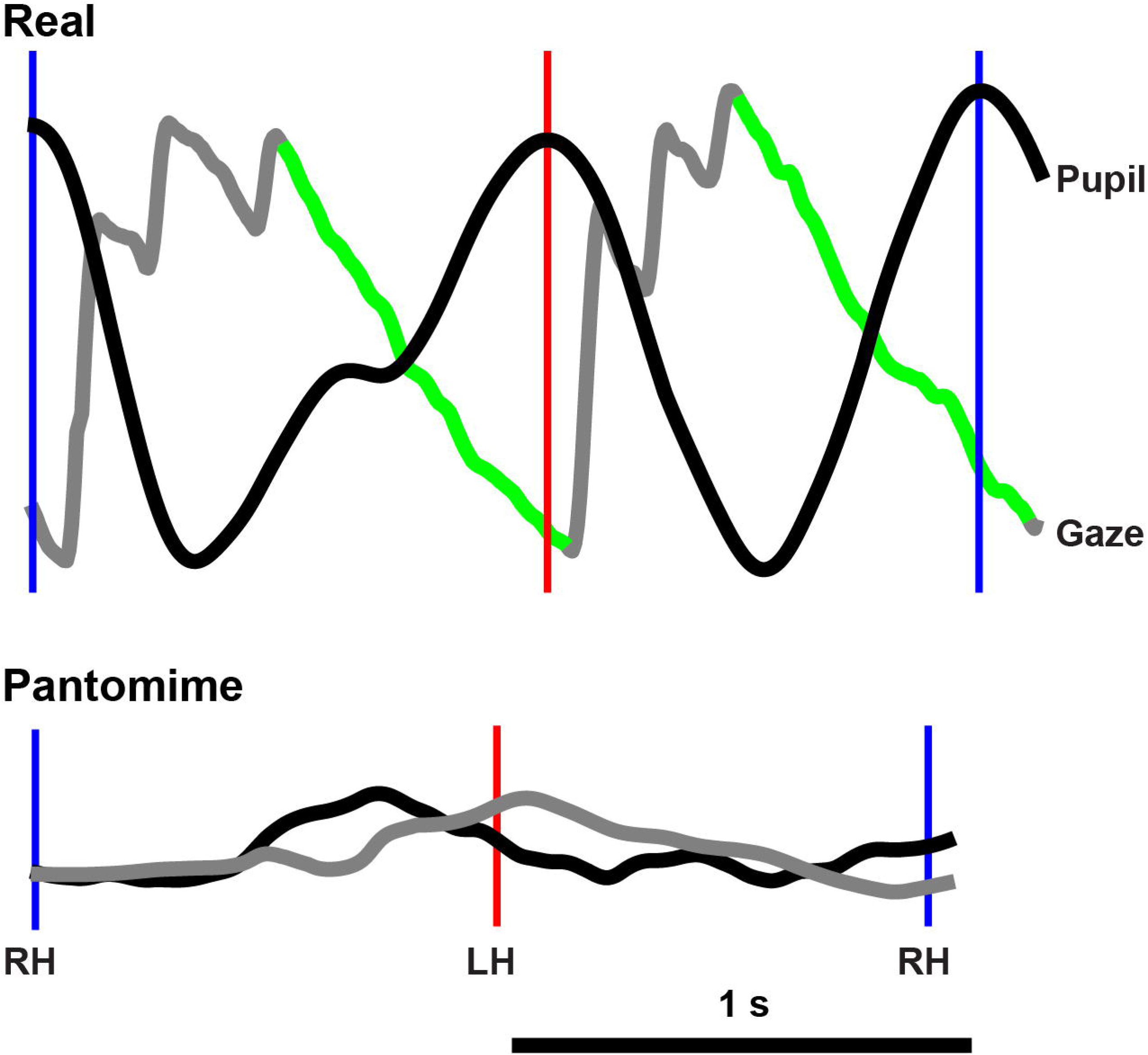
Eye movement (gray) and pupil response (black). The upper panel shows the eye movement tracked the cue to which the participants reached (green portion of the eye movement trace) and then disengaged to search for the next cue. Associated with visual tracking, pupil diameter increased to reach a maximum at about the time that the string was grasped. The lower panel shows that there was no associated eye tracking or pupillary dilation associated with pantomime grasping. LH (Left Hand), RH (Right Hand).

#### Visual cue condition

In the visual cue condition, all 10 participants directed their gaze to the marker on the string as the reaching hand made an up movement. They tracked the marker to about the point that the hand grasped the string at the marker, following which gaze was directed upward to the location of the next marker. In addition to visually tracking the marker on the string, the diameter of the pupil increased as the hand approached the cue and decreased as the hand grasped and pulled.

In the pantomime visual cue condition, all of the participant directed their gaze upward, with 4 participants directing their gaze above the location where they pantomimed a grasping movement and 6 participants directing their gaze to roughly the same location that they pantomimed a grasp. Only one of the participants in the visual cue condition displayed gaze tracking movements similar to the cue tracking movements displayed in the real string-pulling condition. There were no changes in pupil diameter in association with hand grasping at the pretend string.

#### Non visual control condition

In the real non visual control condition, of 10 participants, 5 directed their gaze onto the string at point above their grasp point on the string, 3 directed their gaze onto the string at the location that they grasped the string, and 2 directed their gaze to a point below the location on the string that they grasped. Gaze was mainly fixed on the string at about the same location throughout the string-pulling sequence. There were no hand movement related saccades and the diameter of the pupil did not change during string-pulling movements. Thus, although the participants did look at the string as they were pulling, they did not reliably look at a point on the string at which they grasped. When asked where he was looking, a participant who looked well above the grasping point on the string relied, “It is like driving a car, you look way down the highway not directly in front of you.”

#### Visual attention

For the string-pulling groups that did not wear eye tracking glasses, gaze orientation as assessed by head angle was similar to that obtained in the eye-tracker experiment. A correlation between the head orientation scale and the grasp location scale gave a significant relation when the participants reached to a visual cue (r = 1) or pantomimed the same movement (r=0.981). In this version of the string-pulling task, the participants always looked up and reached above their head (as is also reflected in the frequency and amplitude of their reach movements, see below). There was a significant correlation between head orientation and grasp location when the participants reached for a real string when blindfolded (r = 0.603) or pantomimed the movement (r = 0.680), except that in this task the participants all looked straight ahead, and their grasps were mostly made at the level of the face. There was no significant correlations between head orientation and grasp location in other variations of the real or pantomime conditions; control unmarked string, real (r = 0.0329), pantomime (r = −0.094); control marked string real (r = 0.192), pantomime (r = 0.0242).

### Hand Movements Real and Pantomime Grasp and Release

Figure 3 gives an example of hand movements for release and grasp in the real and the pantomime string-pulling conditions. The initiation of the movements (grasp or release) is shown in the A-panels, the midpoint is shown in the B-panels, and the completion shown in the C-panels. Figure 4 summarized the rating scores of the release and grasp movements, based on a 3-point rating scale (finger opening or closing in the sequence of fingers 5 through 1 = “0”, concurrent opening and closing of all fingers = “0.5” score, and finger opening or closing in the sequence of fingers 1 through 5 = “1” score).

**Figure 3.**
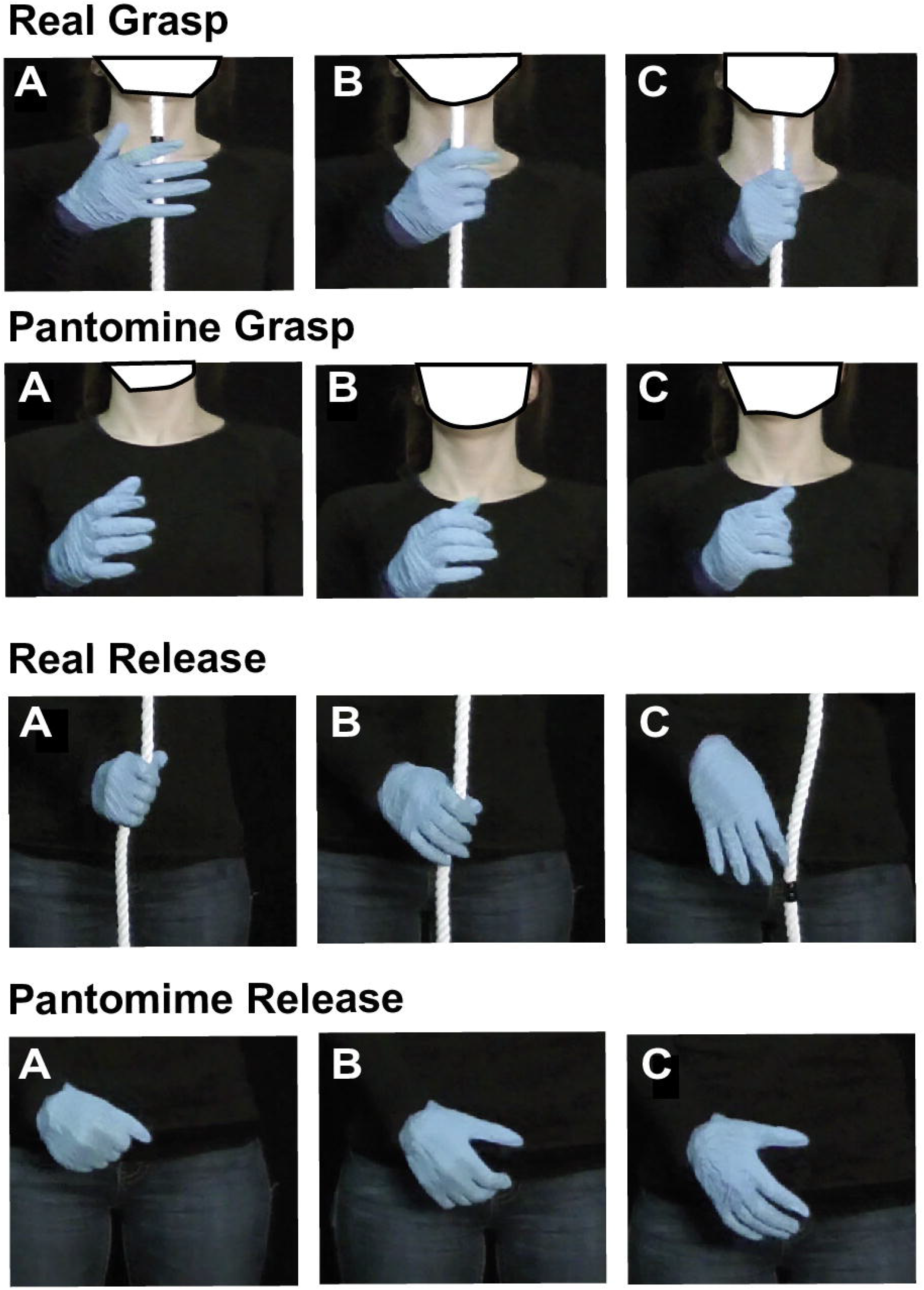
Hand shaping movements to grasp and release the string in real string-pulling and pantomime string-pulling. A, the point of movement initiation, B, midpoint of hand shaping to grasp and release, C, the point the string is released. Note: (1). to make a real grasp of the string the fingers are closed in the order 1 through 5 (pinki first thumb last) and to make a real release the string the digits are opened in the order 1 through 5. (2). To make a pantomime grasp, the fingers are closed in the order 5 through 1, and release the string the fingers are opened in the order 5 through 1.

**Figure 4.**
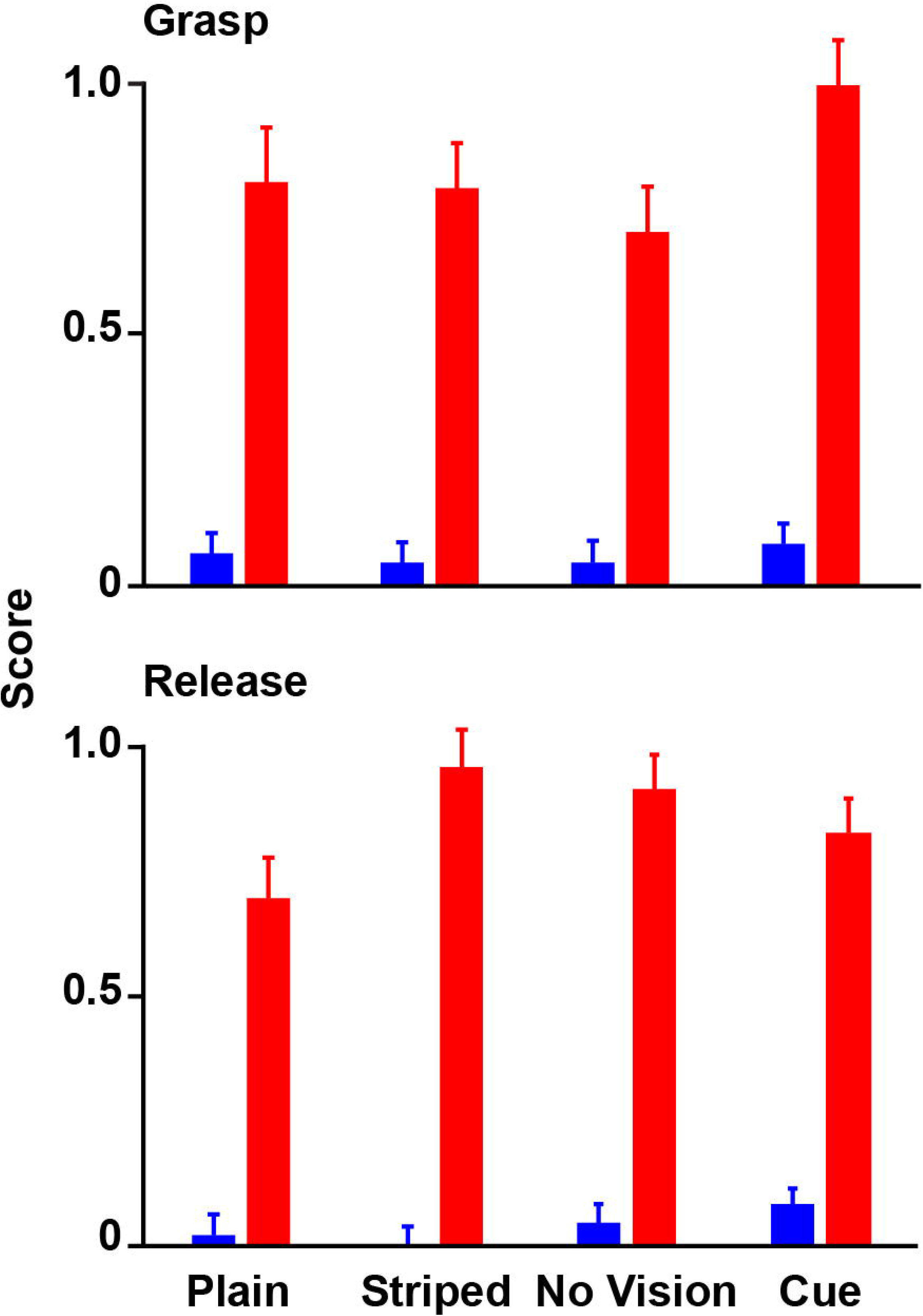
Scores (mean±se) for real and pantomime string release and for real and pantomime grasp. Scores were “0”, the fingers open or close in the order 5 through 1, “0.5”, the fingers open or close concurrently, “1”, the fingers open or close in the order 1 through 5. Note: the 5 through 1 order of finger movement of real-string pulling is reversed to 1 through 5 for pantomime string-pulling.

#### Grasp

To grasp a real string, the hand adducts so that the palm of the hand is aligned with the string, which is in turn aligned with the body midline. The fingers fully open and extend before the grasp and then fingers 5 and 4 usually make first contact with the string in order to gather the string toward the palm. As the hand begins to make a downward movement, the digits close in a sequence of 5 through 1 to fully grasp the string. The grasp for the pantomime string pulling is different. First, as the hand approaches the “string”, the digits do not fully extend or open. Second, the fingers usually close in the reverse order, 1 through 5. Third, the hand closing sometimes occurs in place and sometimes continues its upward movement as the grasp takes place. There was a significant difference between the scores for grasp of the string between the real and pantomime conditions, F(1,40)=223.23, p<0.000, with the 5-1 sequence used for real string-pulling and the 1-5 sequence used for pantomime string-pulling. This difference occurred for all of the groups for both sexes, as the group and sex effects were not different, nor were there significant interactions.

#### Release

To release the string in the real condition, the hand pronates so that the palm is directed toward the body, the fingers extend downward as they open, and they open in the sequence 5 through 1, with fingers 1 and 2 being the last to release the string. When the string is fully released the fingers are fully extended with the palm facing inward toward the body and the digits directed downward. The release for pantomime string-pulling is different. First, the fingers extend only partially in opening to release the “string”. Second, the fingers open in the order, 5 through 1. Third, the fingers may not extend downward and hand opening often begins as the hand starts to make an upward movement. There was a significant difference between the scores for release of the string between the real and pantomime conditions, F(1,40)=357.8, p<0.000, with the 5-1 sequence used for real string-pulling and the 1-5 sequence used for pantomime string-pulling. This difference occurred for all of the groups for both sexes, as the group and sex effects were not effects were not significant, nor were there significant interactions.

### The timing of hand opening to grasp

Figure 5 shows a representative y-trajectory of a real string-pulling cycle (top) and a representative y-trajectory of a pantomime string-pulling cycle (bottom). The dotted lines superimposed on the trajectories represent the point of maximum hand opening relative to peak amplitude associated with the upward movement of the hand. Inspection of the video of hand opening for the real groups indicated that they opened the hand fully as the string was released, displayed slight flexing and closing of the digits as the hand reached the midpoint of the torso, and displayed greater opening and extension of the digits just before the grasp. Participants in the pantomime groups displayed a gradual opening of the hand as it was lifted to reach a maximum at the time of the grasp. It was not possible to quantify the details or hand rotation on the upward movement from the single anterior view of the video record. An ANOVA on the time to maximum opening before the apex of the up movement indicated that there was a significant difference between real and pantomime reaches, F(1,40)=7.81, p<0.01, but no group, sex, or interaction differences.

**Figure 5.**
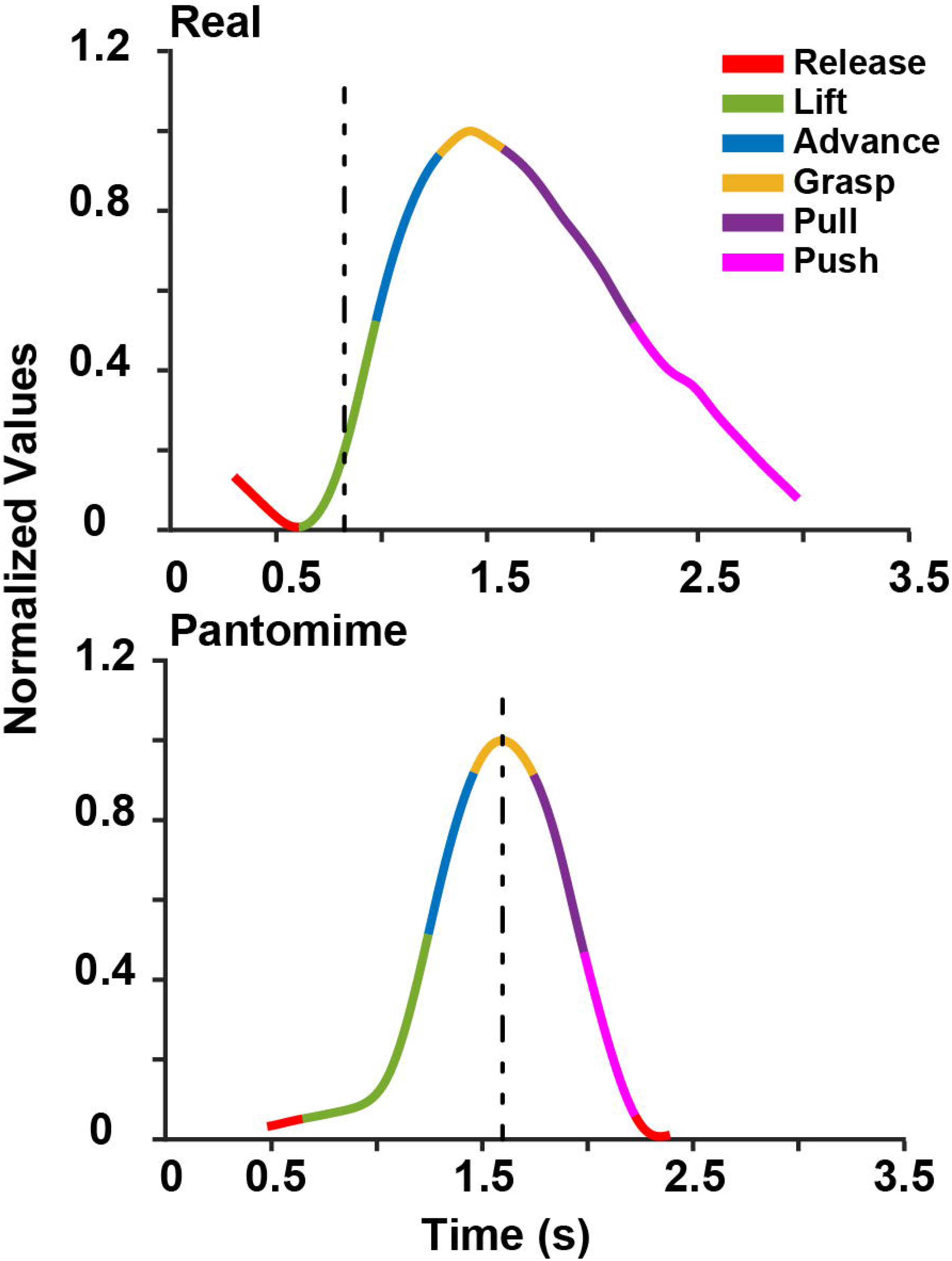
A representative Cartesian plot of hand movement in the y-direction (up/down) for a real string-pull (top) and a pantomime string-pull (bottom). The vertical lines represent the point prior to peak amplitude that the hand fully opens after release. Note: for real string-pulling, the hand fully opens at the point of initiation of the lift, whereas for pantomime string-pulling, the hand opens fully at about the point that grasping is initiated.

### Real and pantomime velocity profiles

Figure 6 shows a representative velocity profile of a single real string-pulling cycle (top), a pantomime string-pulling cycle (middle), and group values (bottom). The velocity profile of the real string-pulling cycle features two high velocity peaks, a high velocity peak for the up movement and a lower velocity peak for the down movement. Both peaks occurred at about the point that the hand passes the midpoint of the torso and at this point the two hands were juxtaposed at the midpoint of the torso. The velocity profile a the real reach also features two points of minimum velocity, the first as the fingers close to grasp the string and the second as the fingers open to release the string, and the minimums occurred sequentially, the release by one hand following the grasp by the other hand.

**Figure 6.**
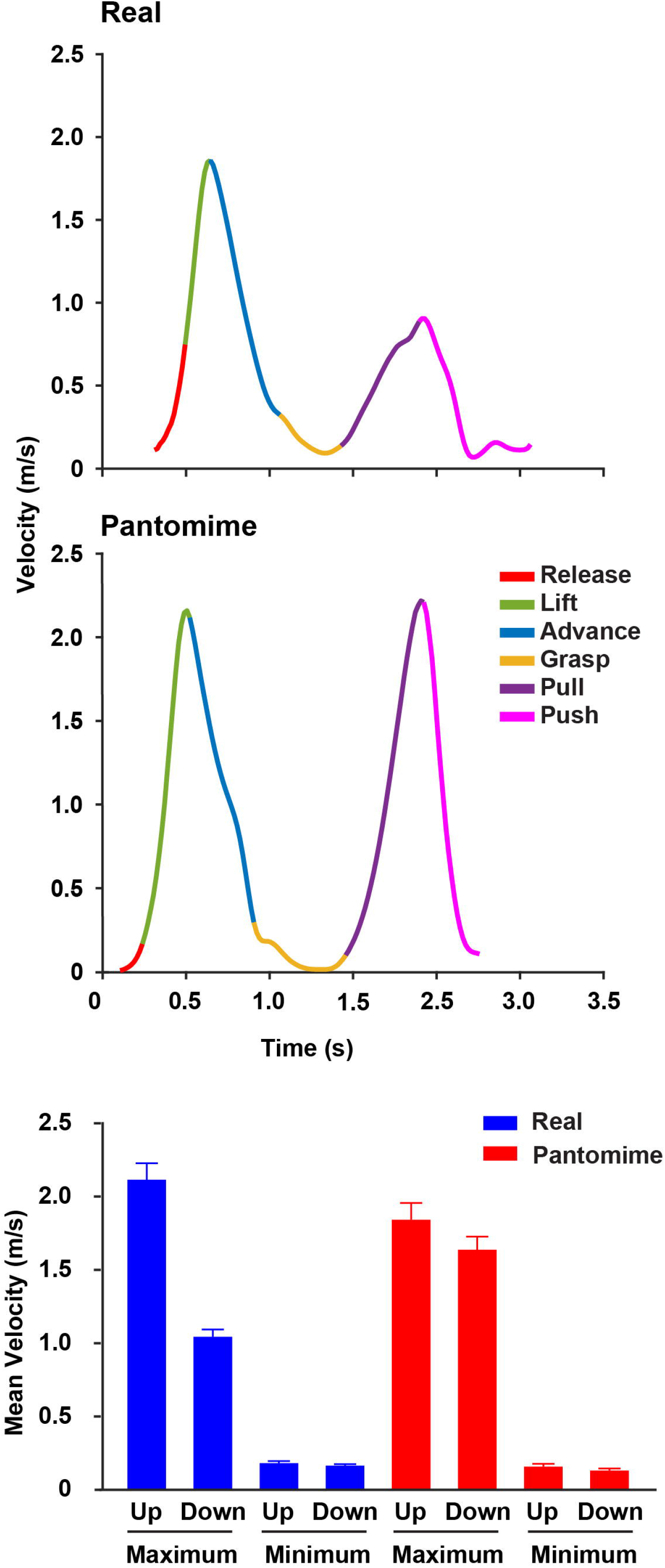
Representative velocity profiles for a real string pull (top) and a pantomime string pull (middle) and group values (mean±se) for real and pantomime maximum and minimum velocity. Note: (1) Maximum velocity occurs at the midpoint of the torso at about the transition of the lift and advance movements; (2) The maximum velocity of the down movement is lower than the maximum velocity of the up movement for real string-pulling, whereas up and down velocity is equivalent for pantomime string-pulling; (3) Minimum velocity occurs at the grasp point for the up movement and at the release point for the down movement for both real and pantomime string-pulling.

The profile of pantomime string-pulling features high velocity peaks for upward and downward movement that are equivalent, and both occur as the hands are juxtaposed passing through the torso midpoint. The pantomime velocity profiles also feature two minimums, one at the grasp and one at the pantomime release, the minimums frequently feature a pause in motion, and the release by one hand and the grasp by the other occur almost simultaneously.

The analyses of the maximum and minimum velocities for the up and down movements of real and pantomime reaches gave a significant difference in maximum/minimum velocity for the comparison of real vs pantomime conditions, condition F(1,40)=418.58, p,0.000, and a significant difference in the measures for the direction of hand movements, direction F(1,40)=102.884, p<0.000. There was a significant interaction between maximum/minimum velocity and direction, F(1,40)=90.992, p<0.000, between real vs pantomime and direction, F(1,40)=98.956, p<0.000, as well as a significant three way interaction between maximum/minimum, direction and condition, F(1,40)=118.69, p<.000.There were no differences in the maximum, and minimum velocities as function of group or sex, and none of the interactions featuring group or sex were significant.

### Frequency and amplitude in real and pantomime string-pulling

Figure 7 gives a summary of the scores for frequency, number of cycles per/sec, and amplitude, apex to apogee of the upward/downward distance, of string pulls for the real and pantomime conditions.

**Figure 7.**
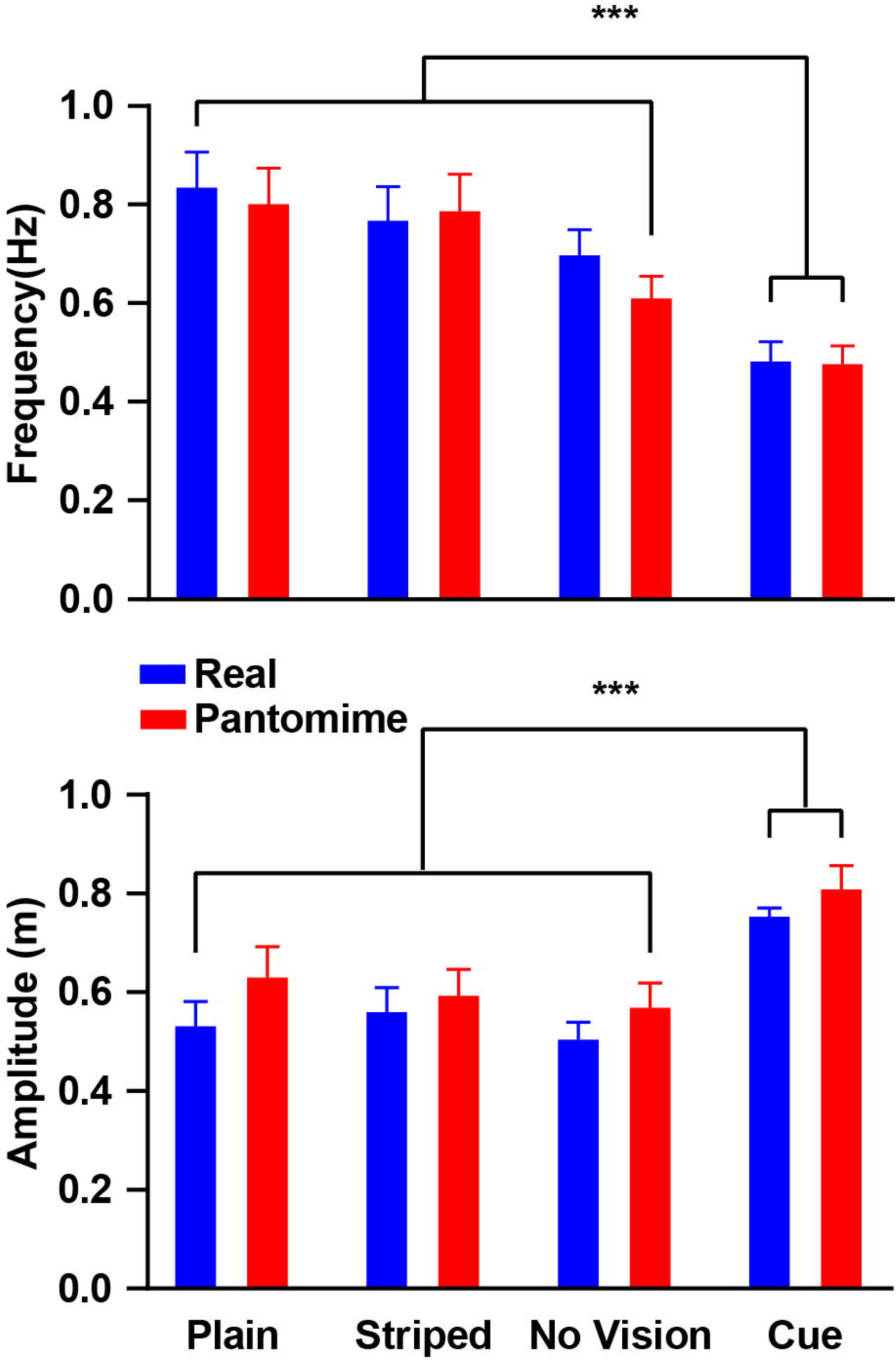
Frequency (top) and amplitude (bottom) (mean±se) for real and pantomime string pulling. Note: (1) Frequency and amplitude do not differ in real and pantomime string-pulling; (2) Frequency is lower and amplitude is higher for both real string-pulling and pantomime string-pulling in the vision task in which participants were required to grasp the string at markers to 30 cm intervals.

#### Frequency

A summary of string-pulling frequency is displayed in Figure 7A. There was a significant group effect, F(3,40)=9.16, p<.000. Bonferroni follow-up tests indicated frequency by the group in the vision-cue condition was slower than in the other groups, which did not differ from each other (vision-cue vs no vision, p=0.000, vision-cue vs control plain, p=0.002, vision - cue vs control striped, p<0.001). The comparison between string pulling frequencies for the real vs pantomime conditions gave no significant difference, condition F(3,40)=0.41, p=0.24. There was no significant sex difference or significant interaction between group and sex.

#### Amplitude

A summary of string-pulling amplitude is displayed in Figure 6B. There was a a significant group effect, F(3,40)=8.857, p<0.001. Bonferroni follow-up tests indicated that string-pulling amplitude by the group in the vision-cue condition was higher than that of the other groups (vision-cue vs no vision, p=0.003, vision-cue vs strip=0.006, vision-cue vs plain, p=0.033), which did not differ from each other The comparison between string-pulling amplitudes also gave a significant difference, with the amplitude of reaches in the pantomime condition being slightly higher than the amplitude for reaches in the respective real conditions for all groups, condition F(1,40)=13.71, p>0.001. The difference may have been due the hand lowering that occurred for real string-pulling as the string was grasped vs the hand raising that often occurred for pantomime string pulling as the string was grasped. There was a significant sex difference F(1,40)=10.179, p=0.003, in which the amplitude for the male participants was higher than that for the female participants. This difference is not surprising as females were shorter than the males. None of the interactions involving hand and sex were significant.

#### Independent and Synchronized Up/Down Movements of Real and Pantomime String-Pulling

Figure 8 shows velocity profiles of representative right-hand (solid line) and left-hand (dotted line) string-pulling cycle of a participant making a real string-pulling movement (top), a pantomime string-pulling cycle (middle), and the group values for combined movements made in both test conditions (bottom). The dotted vertical lines superimposed on the velocity profiles show the period of the string-pulling movement during which both hands pulled together (synchronized hand movements) and for the rest of the movements the hands were making movements in different directions (independent hand movements). For most of the string-pulling movement in the real and pantomime test conditions, the hands were engaged in making independent directional movements – one hand going up and the other going down and one hand releasing and the other grasping. For a portion of the act, however, there were combined movements during which both hands were going down at the same time. The portion of the-string pulling sequence involving both hands that is demarked by the dotted lines in Figure 7 indicates the portion of combined movement.

**Figure 8.**
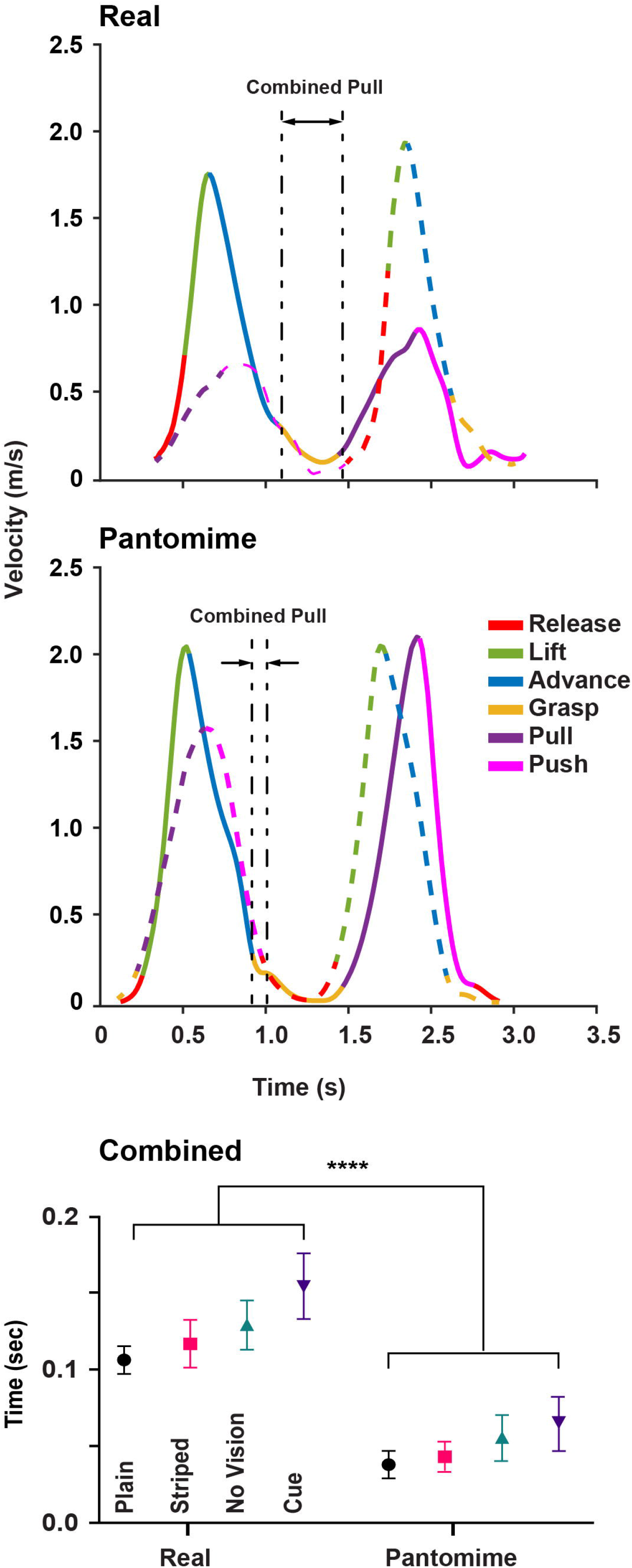
Representative right (sold line) and left (dotted line) velocity curves for real (top) and pantomime (middle) string-pulling and values (mean±se) for concurrent movements of the right and left hands. The period between the vertical lines represent the time that the two hands are moving in concurrently (downward as both advance the string). Note: (1) For real string-pulling the concurrent movement occurs as the right-hand grasps and pulls and the left-hand pushes; (2) For pantomime string pulling the concurrent movement is shorter and occurs as the right-hand grasps and the left-hand releases the string; (3) Concurrent hand movements are longer for all groups engaged in real string-pulling than in the corresponding pantomime groups.

For real string pulling, the hands moved together when one hand was grasping and beginning the pull and the other hand was making the push. The difference between real and pantomime string-pulling was that the combined movement duration was much longer for real string-pulling than it was for pantomime string-pulling. The group measures showed that the total time making combined movements for the real string-pulling test condition was significantly longer than that of the pantomime condition, Condition F(1,40)=86.155, p<0.000. In addition, there was an overall group effect, Group F(3,40)=2.955, 0.045. The combined durations were slightly longer for the participants in real and pantomime cued group than for the other groups. This was likely related the overall longer duration string-pulling movements made by participants in the cued group. There was no effect of sex.

### Bimanual coordination of Left and Right Hands in Real and Pantomime

Figure 9 (top, left) shows the relationship between a complete cycle of a series of left hand string-pulling movements and right hand string-pulling movement for real and pantomime string pulling. For real condition there was a slight lag between the left and right hand movements but for the pantomime conditions, movements were synchronous, when the left hand was going up the right hand was going down. The raw y-data from the left and right hands were used to evaluate the correlation of the hands (Figure 8, right). Repeated measures ANOVA conducted on initial correlation revealed a significant main effect of condition, F(1, 44)=19.308, p<0.000, but no significant effect of group or group by condition interaction.

**Figure 9.**
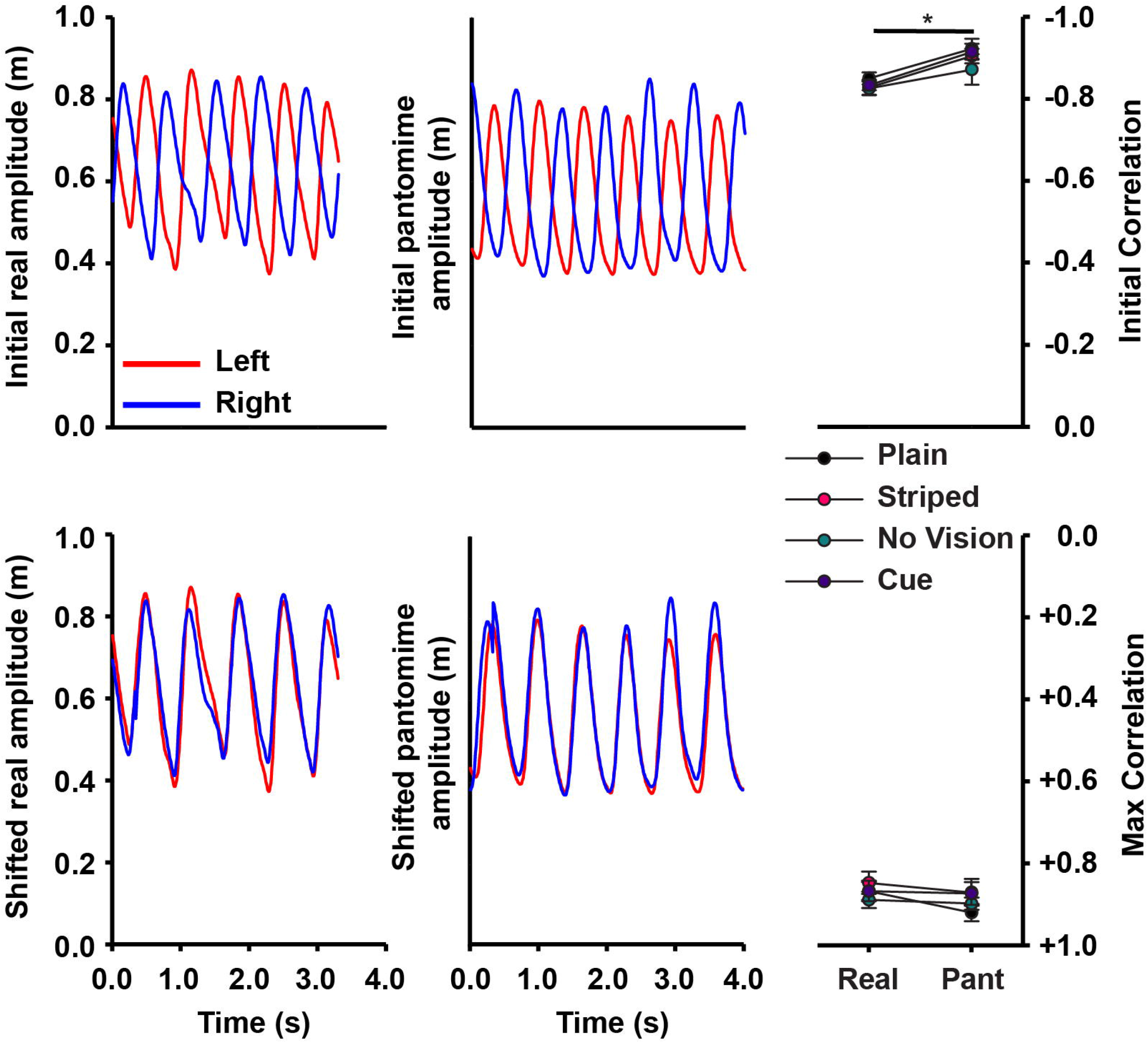
Relationship between left and right hand string-pulling movements. A. Change in y-values for left and right hand movements for sequences of string-pulling for real and pantomime string-pulling. B. Correlations between left and right movements provide a greater negative correlation for pantomime than real movements. C. Change in y-values for real and pantomime reaches are shifted to the position of maximum correlation. D. Correlations between left and right hand movements are highly positive, showing that for both real and pantomime movements, the left and right hands are making very similar movements in string-pulling.

To asses whether the movements of the right and left hands were equivalent, the raw y-data for the right hand was shifted frame-by-frame until the maximum correlation was achieved between the left and right raw y-data for each participant (see Figure 8, bottom, left). Repeated measures ANOVA conducted on maximum correlation (Figure 8, bottom, right) failed to reveal a significant main effect of group, F(3, 42)=1.310, p=0.284, condition, F(1, 42)=1.360, p=0.250, or condition by group interaction, F(3, 42)=0.384, p=0.765. Thus, the left and the right hands are making the same, but reciprocal movements.

### Asymmetry and Spatial Occupancy in Real and Pantomime String-Pulling

Figure 10 gives a Cartesian representation of the topography and velocity (indicated by color coding) of string-pulling movements made by a representative subject engaged in real string-pulling (top) and pantomime string-pulling (bottom). The cartoons of the left- and right-hand movements illustrate the spatial occupancy of the movements and the degree to which the movements of the two hands are symmetric. For this participant, the movements of the right and the left hand are symmetric as they were for other participants in the real string-pulling conditions. The movements of pantomime are quite asymmetric, as they were for other participants in the pantomime conditions, with one hand having a higher rise time and greater spatial occupancy than the other hand. This asymmetry was not lateralized, however.

**Figure 10.**
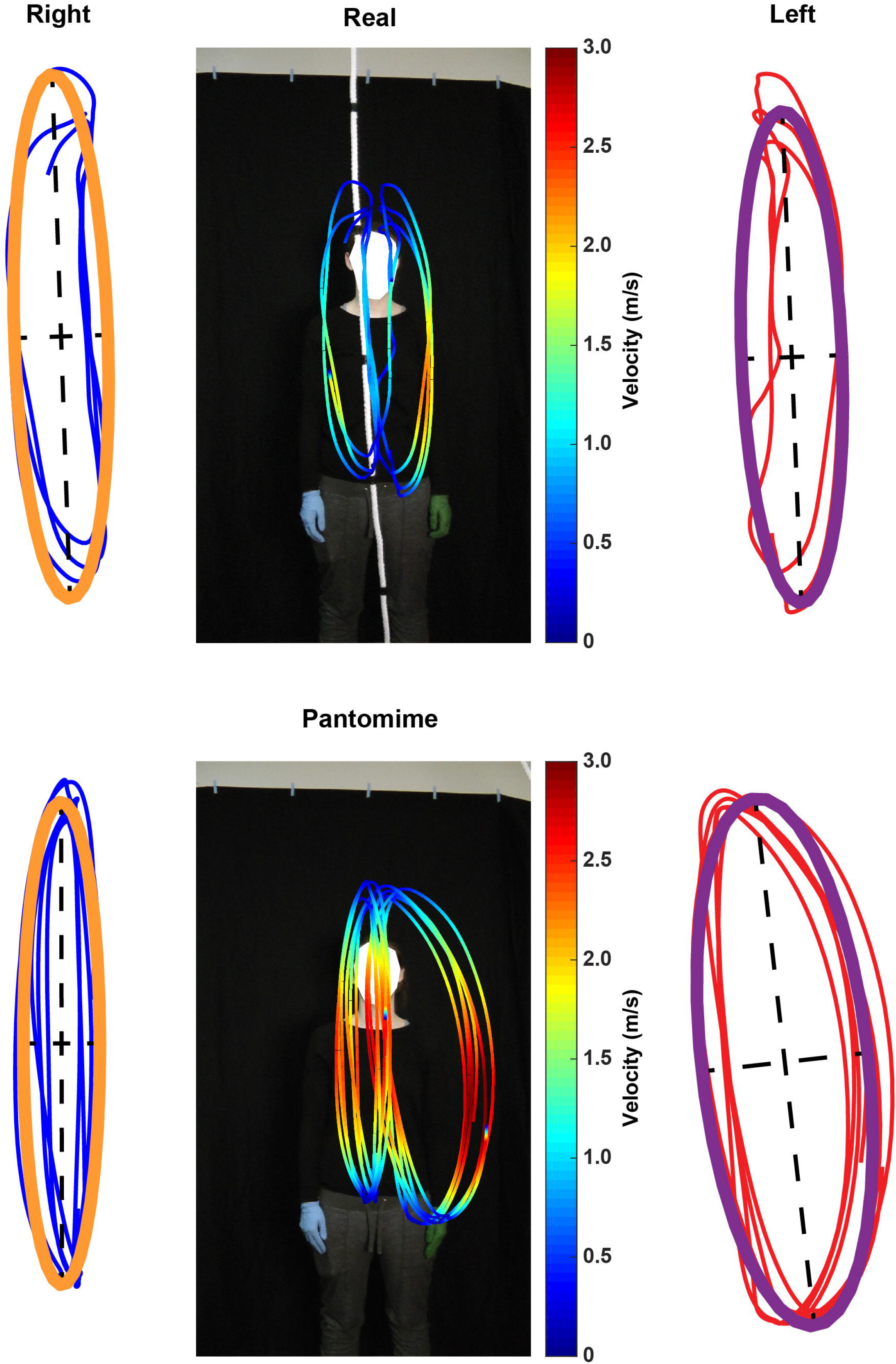
Representative Cartesian trajectories of right- and left-hand real sting-pulling (top) and pantomime string pulling (bottom). In the trajectory plots of the hand movements, the thin (red and blue) lines represent each cycle of movement and the thicker (purple and golden) lines represent overall elliptical fit. Note: the spatial similarity of the real-string pulling vs dissimilarity of pantomime-string pulling.

Figure 11 summarizes group values for the parameters of spatial occupancy of the right-hand and the left-hand movements for real string-pulling and pantomime string-pulling:

**Figure 11.**
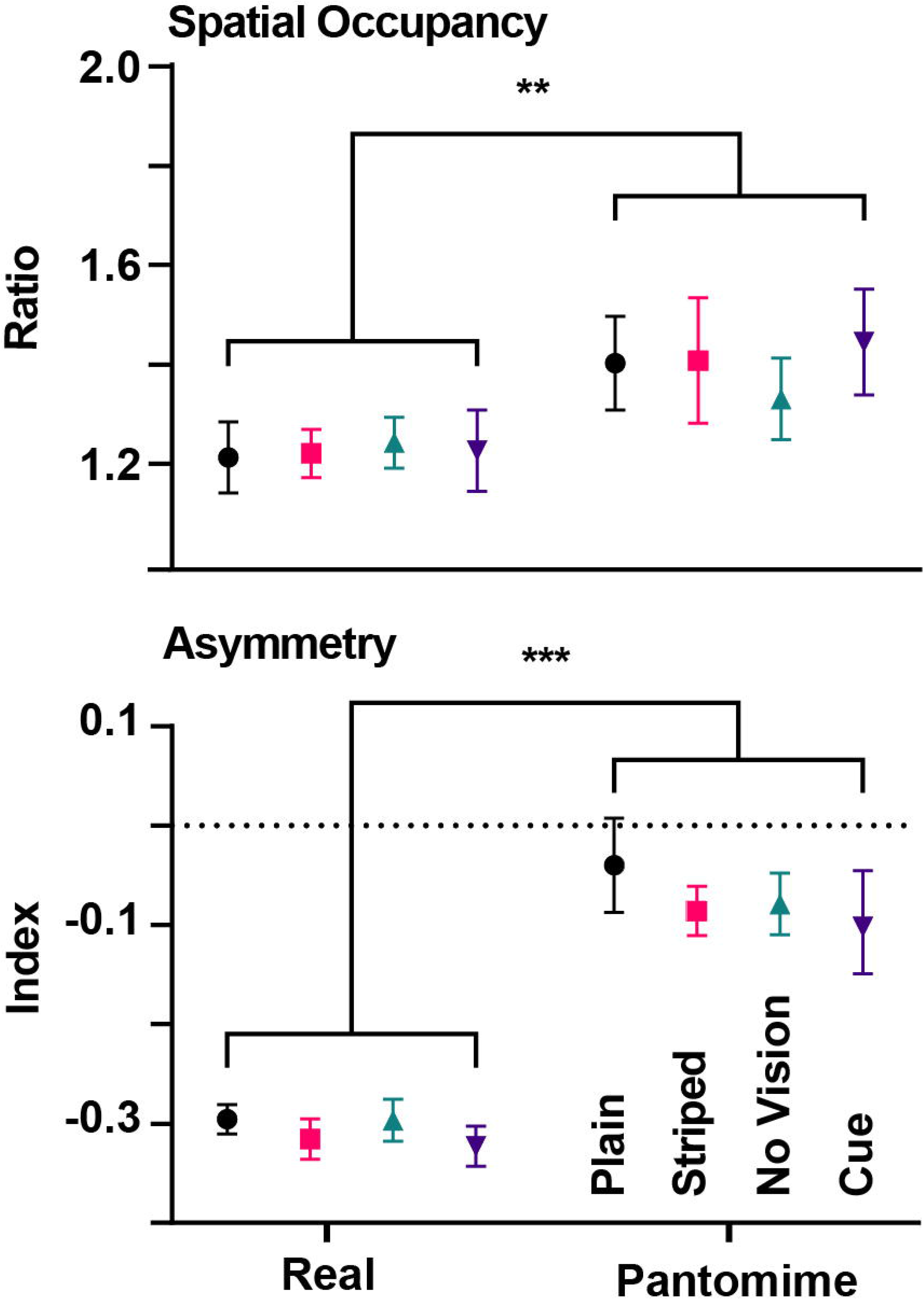
Spatial occupancy (top) and asymmetry (bottom) of trajectory values (mean±se) for real and pantomime string-pulling. Note: (1) Spatial occupancy (trajectories are more expansive) of pantomime than for real string-pulling; (2) Trajectories are more asymmetric for pantomime than for real string pulling.

#### Spatial occupancy

Figure 11A gives values for spatial occupancy. The ANOVA showed that there was no hand effect with respect to spatial occupancy, hand F(1,40)=0.104, p=0.749, so for some subjects, spatial occupancy was just as likely to be larger for the right hand as for the left hand. There was an effect of test condition, in which the spatial occupancy of real string-pulling was smaller than the spatial occupancy of pantomime string-pulling, condition F(1,40)=4.929, p<0.000. Thus, the presence of the string constrained the hands to more up/down movement than side/side movement. There were also effects of group, F(3,40)=10.125, p<0.000, and sex, F(1,40)=10.025,p=0.003, and there were no significant interaction effects. In general, the spatial occupancy results for group and sex mirrored those of amplitude (above), in which the larger amplitude of pantomime hand movements were associated with larger spatial occupancy as were the larger amplitude movements of participants pulling in the vision-cue test condition. Accordingly, the smaller amplitude movements of female participants are associated with smaller spatial occupancy movements, reflecting body size.

#### Asymmetry Index

Figure 11B shows the values for the asymmetry index, the difference in space occupied by the movement of one hand vs the space occupied by the other hand. Thus, for real string pulling the excursions of the two hands were quite similar, whereas for pantomime the excursions of the two hands were different, without evincing a bias for one hand or the other. An ANOVA confirmed this difference, giving a significantly lower index for real string-pulling compared to pantomime string-pulling, test F(1,40)=103.517, p<0.000. There were no significant differences as a function of group or sex and there were no significant interactions.

## Discussion

Frame-by-frame video inspection, eye tracking and MATLAB based kinematic monitoring describe the sensory control, movement, and kinematics of real hand-over-hand string-pulling and pantomime of the movement. Sensory control varied as a function of the string-pulling task, with participants in a cued task visually tracking the cue, in a noncued task monitoring the string but not grasping location, and in a nonvision task relying on somatosensory cues. Pantomime string-pulling movements reflected features of real string-pulling but also featured systematic changes in hand movement and temporal coupling of movement. This description adds to the description of string-pulling behavior described for other animal species. The study also confirms a contemporary literature showing differences between real and pantomime action. The results support theories that real and pantomime movements may have different neural substrates but the uniformity of some features of pantomime movements suggest that part of pantomime movement may be the use of gestures such as those related to manual communication.

A surprising finding is that direct visual control hand placement on the string did not seem essential for performance. Many studies that describe reaching for target objects or food items or pointing toward a target report that gaze is fixed or anchored on the target to which the hand is directed (Prablanc et al., 1979; Neggers and Bekkering, 2000, 2001; de Bruin et al., 2008; Sacrey and Whishaw, 2012). Here gaze anchoring was usually absent, although participants did look at the string. Eye tracking and measures of head orientation showed that when participants did look at the string, they often did so at a point above or below their hand contact points. That visual guidance was not essential was confirmed by identical kinematic results obtained when participants were blindfolded. With participants instructed to grasp cues on the string, gazed was directed to the cues, the cue was visually tracked, and the pupils dilated at about the point of the grasp. In the absence of visual control, it seems likely effective string pulling occurred because the participants always had one hand on the string, thus providing some information about its location and because the string was typically located on the body midline, providing further information about location. In addition, inspection of the participants’ grasp indicated that they approached the string with a fully open hand and used their distal fingers to contact/gather the string before grasping. Previous work shows that when participants reach without foviating a target, e.g, reaching when blindfolded (Karl et al., 2012) or into peripheral vision (Hall et al., 2014), they use an open hand to find the target and then use touch to guide their grasp. It is interesting that in the many studies of string-pulling in many animal species only one study featuring mice, who monitor string location with their vibrissae, has previously analyzed string-pulling sensory control (Blackwell et al., 2018a).

Finger movements for real vs pantomime movements featured opposite sequences. For real grasping, grasping the string used fingers 5 through 1 as did release of the string. For the pantomime, finger sequence was usually reversed, 1 through 5. Inspection of a grasp showed that the string was usually gathered with fingers 5 and 4, following which the other fingers successively closed. To release the string, the hand and digits were extended laterally away from the string leaving fingers 1 and 2 as the last fingers to release. Despite the fact that the participants making a real string-pulling grasp used a whole hand grasp for purchase, their hand made an arpeggio approach to the target similar to that described for participants grasping a static object such as a food item, except that the precision movements of purchase generally involves only the first two or three digits making contact on an item (Sacrey et al., 2009). An arpeggio movement is featured in many forelimb acts, including placing a hand when crawling or a foot when walking, suggesting that it is a basic pattern(Whishaw et al., 2010). It is surprising that finger grasping patterns were reversed in pantomime because previous studies of pantomime report the kinematics of grasps are only somewhat altered (Goodale et al., 1994; Kuntz and Whishaw, 2016), and not reversed as is found here. Thus, the change in pantomime grasp and release described here are not simply inaccurate versions of real grasping but rather are quite different movements.

Although the two hands made alternating movements at the same speed to advance the string, the up/down movements of pantomime were different from the up/down movements of real string-pulling. For real string-pulling, the down hand movement was slow and the up movement was fast, whereas for pantomime both hands moved at the same speed. In some respects, this asynchrony of real string pulling is similar to that of a step in which the swing phase is rapid relative to the stance phase (Perry, 1992). The synchrony observed in pantomime string pulling may in part derive from the fact that pantomime is free from the constraints imposed by grasping and releasing the string and in part from the emergence of a simplified pattern of limb alternation. For example, it is suggested that the kinematics of limb movement represent two influences, that of a pattern generator that produces the movement and that of control mechanisms that puts the limb to a specific functional use (Marder and Bucher, 2001; Swinnen, 2002). A pantomime movement may reflect mainly the action of the oscillator, whereas the changes associate with real reaching may reflect the addition influence of a control mechanism. In this respect, it is interesting that a similar simplification of movement is reported for pantomime reach-to-grasp in conditions in which sensory influences are minimized (Kuntz and Whishaw, 2016).

A remarkable feature of pantomime string-pulling was the consistency of the behavior across participants. This consistency raises the question of what degree the pantomime movement is self-copying, the participant is trying to produce the best possible real movement, as opposed to performance of a gesture such as the manual movements accompanying speech. Were pantomime movement self-copying (Goodale et al., 1991; Goodale et al., 1994; Westwood et al., 2000; Milner et al., 2001; Fukui and Inui, 2013; Holmes et al., 2013; Kuntz and Whishaw, 2016), it might be expected that some participants would producing a more realistic movement representation than others. Rather than displaying variability, however, the participants were consistent in reversing the finger sequence and in using a simple up/down movement to represent the string-pulling action. People commonly use a variety of more standard gestures to represent grasping and arm motion actions and the gestures usually consist of representative movements rather than exact action imitations of (McNeill, 1992; Kendon, 2004). The gestures themselves, however, may be influenced by the task at hand, such as the looking up when reaching for a cue on the string. One theory of hand use proposes that gaze anchoring, the reach, and the grasp are each produced by separate parietal cortex to motor cortex pathways, with gaze and the reach mediated by more dorsal pathways and the grasp mediated by a more ventral pathway (Jeannerod, 1981; Jeannerod et al., 1994; Karl and Whishaw, 2013). In bimanual string-pulling the regulation of real and pantomime movement may involve more complex cortical networks than single handed movement (Debaere et al., 2003) but nevertheless, the substitution of gestures for real movement may be part of a perceptual network.

This comparison of string-pulling provides a detailed description of a bilateral movement in which the two hands engage in sequential cooperative movements. The study shows that each facet of string-pulling including visual tracking, grasp/release of the string, the amplitude and frequency of arm movements, and the temporal relations between the two limbs, change when participants pantomime. These human string-pulling movements have some similarity to those displayed by other animals such as rodents, although there are presently no reports that nonhuman animals can pantomime similar movements. The real and pantomime string-pulling task could be used to investigate the neural basis of bilaterally coordinated movements and also as a therapeutic tool in diagnosing brain injury and enhancing recovery from brain injury.

## Author contributions

S.S. and I.Q.W. conceived and designed the study, S.S., K.B. and A.M. performed experimental work, S.S., I.Q.W, A.A.B. and K.B. performed data analysis, S.S., A.A.B., M.H.M., D.G.W. and I.Q.W. wrote the manuscript, which all authors commented on and edited.

## Acknowledgements

S.S. and I.Q.W. will like to thank Hardeep Ryait, for his help in setting-up Pupil-Lab eye tracker.

---

Video 1. Representative video of real string-pulling.

Video 2. Representative video of pantomime sting-pulling.

